# Proteomic analysis reveals regional sex differences in healthy and fibrotic human lung

**DOI:** 10.64898/2026.05.15.725416

**Authors:** Rachel Blomberg, Jeremy A. Herrera, Haley Noelle, Mikala C. Mueller, Maxwell C. McCabe, David A. Schwartz, Chelsea M. Magin

**Affiliations:** Department of Biomedical Engineering, University of Colorado, Denver | Anschutz, Aurora, CO, USA; Division of Pulmonary, Allergy, and Critical Care Medicine, Department of Medicine, University of Colorado, Anschutz, Aurora, CO, USA; Department of Biochemistry and Molecular Genetics, School of Medicine, University of Colorado, Anschutz, Aurora, CO, USA; Department of Microbiology and Immunology, University of Colorado Anschutz, Aurora, CO, USA; Department of Pediatrics, University of Colorado, Anschutz, Aurora, CO, USA

## Abstract

Biological sex has systemic effects on gene expression, cell behavior, and disease etiology. Despite these widespread effects, sex as a biological variable is understudied, particularly in chronic lung diseases. In idiopathic pulmonary fibrosis (IPF), 70% of patients are male, and male patients have overall worse survival post-diagnosis. While behavioral differences between sexes might account for some of the epidemiological differences, the contribution of underlying biology is not known. In this study, we performed regional proteomic analysis via laser-captured microdissection-coupled mass spectrometry and analyzed the data for sex-biased protein expression. We discovered that even in control lung, sex differences existed in both airway and alveolar regions. Sex differences became more pronounced in diseased regions, with sex-biased expression of diverse proteins including those involved in extracellular vesicle secretion, cellular metabolism, and extracellular matrix remodeling. These data suggest that baseline sex differences in lung proteome may contribute to sex-specific susceptibility, progression, and clinical outcomes in IPF, underscoring the need for future mechanistic and clinical studies to account for sex as a biological variable.

## Introduction

Across a variety of biomedical research fields, the importance of sex as a biological variable is becoming increasingly appreciated (1). Despite growing acknowledgement that biological sex can impact human health and disease, and even policies encouraging the use and reporting of both sexes in federally funded research, mechanistic research into these effects still lags behind (2, 3). What we do know is that sex differences exist across the body, even in tissues and organ systems that are not classically thought of as having sexual characteristics. Some of these differences are likely because of systemic sex-hormone signaling, but there are also significant differences in overall gene expression in organs as diverse as skin, spleen, liver, and lung (4–6).

In the lung, multiple diseases show sex-biased epidemiology, including pulmonary arterial hypertension (PAH), asthma, lung cancer, chronic obstructive pulmonary disease (COPD), and idiopathic pulmonary fibrosis (IPF) (7–12). Historically, prevalence of chronic lung disease was higher in men than in women, and attributed to behavioral and occupational differences, most particularly higher rates of smoking in men; however, as rates of smoking in women have increased, and when data analysis is normalized for smoking history, more complex trends have begun to emerge (13).

Recent statistics demonstrate a higher prevalence of COPD in women than in men, most notably in younger populations, and even in the face of low tobacco exposure (11, 14). Some potential underlying mechanisms for this discrepancy include complex interplays between hormone signaling pathways, genetic and epigenetic factors, metabolism, and chronic inflammation. Both estrogen and testosterone can have profound effects on pulmonary biology, with estrogen implicated in cellular proliferation and testosterone in alveolar destruction, both of which are hallmarks of COPD (15).

Beyond sex hormones, a recent study identified multiple sex-biased genetic loci involved in COPD risk, though the critical factors and mechanisms underlying these trends are yet to be fully explored (16). On an epigenetic level, sex differences in methylation also potentially contribute to COPD risk, including potential sex-specific regulation of a variety of extracellular matrix (ECM) related genes as well as *CYP1B1* (17, 18). This gene encodes for an enzyme in the cytochrome p450 family that is upregulated in response to smoking and plays roles in both metabolism of xenobiotics (e.g., environmental pollutants) and estrogen signaling (18, 19). Meanwhile, women exhibit high levels of inflammation and oxidative stress in the lung as a response to smoke exposure, a phenotype which might be due to the general pro-inflammatory effects of estrogens, in contrast to more anti-inflammatory effects of testosterone (20, 21).

Many similar risk factors, especially smoking and age, underlie both COPD and IPF, but in contrast to the female-bias in COPD prevalence, IPF is well-documented as a male-biased disease. IPF is a disease characterized by progressive scarring, in which repeated lung injury (e.g. caused by inhalation of smoke/particulate matter) induces an aberrant wound healing response that results in excessive deposition of dense, fibrillar ECM (22). Approximately 70% of IPF patients are men, and men have worse overall survival after diagnosis than women (12, 23, 24). As with COPD, historically higher rates of smoking and occupational exposure in men were considered a predominant force behind these observed sex differences, but statistical interactions between smoking and sex as a risk factors suggest that other biological mechanisms also play a role (25).

Discovering these mechanisms is complicated by the fact that many rodent models fail to recapitulate sex differences seen in humans. The classic rodent IPF model involves intratracheal administration of bleomycin; however, young rodents show increased bleomycin-induced fibrosis in females, not males (26). In contrast, bleomycin administration to aged mice (52-54 weeks) does recapitulate exacerbation of disease in males, demonstrating an interplay of age and sex in fibrotic development (27). Despite limitations in rodent modeling of IPF, evidence from human data and other disease models suggests that estrogen may play a protective role against fibrotic development (28–31). In one study, a chemically-induced lung fibrosis model showed higher TGFβ-signaling pathways and reduced overall survival in male mice (32). Similarly, silica-induced injury of male and female mice showed increased pulmonary fibrosis and expression of profibrotic SPP1 in both male and ovariectomized female mice (33). On a genetic level, sex-specific polymorphisms are observed in IPF, particularly in genes related to mitochondrial function (34). The predominant genetic risk factor for IPF development is mutation of the MUC5B promoter; and while there is no known sex-association of this mutation, estrogen has been observed to suppress MUC5B expression in the context of activated B-cell lymphoma (35). These studies both demonstrate unique sex-driven mechanisms that might underlie the observed pathology of IPF and also highlight the need for further study into unified pathways of sex differences in IPF.

In this study, we used a proteomics approach to characterize regional sex differences in the lungs of male and female, control and IPF patients. We performed laser-captured microdissection-coupled mass spectrometry (LCM-MS) to assess protein content in control airways and alveoli, as well as IPF-associated airways, alveoli, honeycomb cysts, aberrant basaloid epithelial cells, and fibroblastic foci. This cohort also contained patients with both wildtype and gain-of-function MUC5B promoter variants, and analysis of the data with respect to MUC5B, has demonstrated that the promoter variant induces proteomic changes in control alveoli that may predispose towards IPF, as well as causing protein changes in IPF airways (36), Here, we re-analyze the cohort with respect to sex to identify potential sex-specific drivers of IPF risk and progression.

## Methods

### Human lung tissue

All lung specimens met the criteria for IPF diagnosis following current guidelines. Patients provided written consent and samples were approved for research by the Colorado Multiple Institutional Review Board CB F490 (COMIRB 15-1147) and through the Lung Tissue Research Consortium (LTRC). See Table S1 for patient demographics.

Lung tissue samples used in this study were part of a previously published proteomics dataset generated by our group (36). The dataset was reanalyzed here to assess sex-associated differences that were not evaluated in the first publication.

### Laser-captured microdissection-coupled mass spectrometry

LCM-MS analysis was carried out as previously described (36, 37). 5 µm-thick formalin-fixed and paraffin-embedded sections were stained with hematoxylin and eosin to allow for region selection by tissue morphology before microdissection using the Molecular Machinery Instruments (MMI) CellCut system on an Olympus IX63 microscope. Multiple tissue regions were pooled per patient for a total volume of 0.005 –0.010 mm^3^. Samples were resuspended in 50 mM triethylammonium bicarbonate (TEAB) with 5% SDS for 20 min at 95° and then 2 h at 60° before being cooled to room temperature. Urea and DTT were added to the samples for a final mixture of 50 mM TEAB, 5% SDS, 7.5 M urea, and 10 mT DTT and then sheared using a Covaris S220 focused ultrasonicator. Samples were then sequentially treated with 500 mM iodoacetamide and 12% aqueous phosphoric acid before centrifugation at 12000 RPM for 5 min. The supernatant was removed and mixed with 90% methanol in 100 mM TEAB before being passed through a S-Trap micro column to capture proteins while removing detergents and other contaminants. The column was washed 10 times with 90% methanol in 100 mM TEAB and captured protein was digested in-column for 1 h at 47° with 0.8 µg/µL trypsin solution. Proteins were eluted from the column with 50 mM TEAB and sequentially combined with 0.2% aqueous formic acid and 50% aqueous acetonitrile before lyophilization. Samples were desalted using Oligo R3 resin beads and then resuspended at 50 ng/uL in 0.1% formic acid.

### Protein Identification and Quantification

Digested peptides were loaded into autosampler vials and analyzed directly using a NanoElute liquid chromatography system (Bruker, Germany) coupled with a timsTOF SCP mass spectrometer (Bruker, Germany). Peptides were separated on a 75 µm i.d. × 25 cm separation column packed with 1.6 µm C18 beads (IonOpticks) over a 90-min elution gradient. Buffer A was 0.1% FA in water and buffer B was 0.1% FA in acetonitrile. Instrument control and data acquisition were performed using Compass Hystar (version 6.0) with the timsTOF SCP operating in parallel accumulation-serial fragmentation (PASEF) mode under the following settings: mass range 100-1700 m/z, 1/k/0 Start 0.7 V s cm-2 End 1.3 V s cm-2; ramp accumulation times were 166 ms; capillary voltage was 4500 V, dry gas 8.0 L min-1 and dry temp 200°C. The PASEF settings were: 5 MS/MS scans (total cycle time, 1.03 s); charge range 0–5; active exclusion for 0.2 min; scheduling target intensity 20,000; intensity threshold 500; collision-induced dissociation energy 10 eV.

### Proteomics data analysis

Raw MS data was searched using MSFragger via FragPipe v 20.0. Precursor tolerance was set to ±15 ppm and fragment tolerance was set to ±0.08 Da. Data was searched against SwissProt restricted to Homo sapiens with added common contaminants (20,410 total sequences). Enzyme cleavage was set to semi-specific trypsin for all samples. Fixed modifications were set as carbamidomethyl (C). Variable modifications were set as oxidation (M), oxidation (P) (hydroxyproline), Gln->pyro-Glu (N-term Q), and acetyl (Peptide N-term). Results were filtered to 1% false discovery rate (FDR) at the peptide and protein level and exported directly to Excel for downstream analysis.

Data were analyzed using MetaboAnalyst 6.0 (https://www.metaboanalyst.ca/) accessed February 2026 (38, 39). Data was filtered to remove the lowest 5% of protein identifications by mean intensity value and LFQ intensities were log (base 10) transformed. Multivariate analyses including a two-way ANOVA (p=0.1, FDR) and a principal component analysis (PCA) were performed and plotted following filtering and transformation. For univariate analyses, a partial least squares discriminant analysis (PLS-DA) created variable importance in projection (VIP) scores ranking the contribution of each protein to the model, with VIP scores greater than one indicating a significant contribution to sample group separation. Differentially expressed proteins were identified using a fold-change (FC) threshold of 2.0 accompanied by a raw p-value of 0.1 relative to control samples. In volcano plots, a raw *p*-value threshold of 0.1 was used to identify proteins exhibiting significant sex-associated differences and these proteins were highlighted on the plots. Heatmaps were created using the top differentially expressed proteins as defined by ANOVA (multivariate) or t-test (univariate) and clustered using Ward’s method.

### Pathway analysis

Enrichment analysis was conducted on differentially expressed proteins using Metascape (metascape.org), accessed February 2026 (40). For each gene list, pathway and process enrichment analysis were carried out with the following ontology sources: KEGG Pathway, GO Biological Processes, Reactome Gene Sets, Canonical Pathways, CORUM, WikiPathways, and PANTHER Pathway. All genes in the genome were used as the enrichment background. Terms with a p-value < 0.01, a minimum count of 3, and an enrichment factor > 1.5 were collected and grouped into clusters based on membership similarities. P-values were calculated by Metascape based on the cumulative hypergeometric distribution, and adjusted p-values were calculated using the Benjamini-Hochberg procedure (41, 42). Comparisons of pathway analysis with individual proteins was assisted by the Harmonizome database (https://maayanlab.cloud/Harmonizome) accessed March 2026 (43).

### Immunofluorescent staining

Human lung sections were flash frozen, embedded in OCT, cryosectioned at 10 µm thickness, and stored frozen until staining. Before staining, slides were equilibrated at room temperature for approximately 15 min and then dipped in phosphate-buffered saline (PBS) to clear residual OCT. Slides were lightly fixed in ice-cold acetone for 15 min before being washed in PBS. Lung sections were circled with a hydrophobic pen (Vector Laboratories) and blocked in 5% bovine serum albumin (BSA) in PBS for 30 minutes. Slides were then incubated in primary antibodies or 5% BSA alone (controls) overnight at 4°C. Slides were washed 3x in PBS with 1% Tween20 (PBST) before being incubated for 1 hour in secondary antibodies. Where the antibody panel included a goat-anti-human primary, sections were incubated in donkey-anti-goat secondary alone before washing 3x in PBST and performing a second incubation with any goat secondaries. After the final antibody incubation, slides were washed 2x in PBST and 1x in PBS before incubation for 15 min in 10 µg/ml Hoechst 33342 (ApexBio) in PBS. Slides were washed 3x in PBS and then mounted using Prolong Gold Antifade Reagent (Thermo Fisher). The mount was allowed to cure overnight, protected from light, before imaging on an epifluorescent microscope (Olympus BX63)

### Image analysis

Immunofluorescent images of cryosectioned lung tissue slices were analyzed using Fiji (ImageJ). To reduce background fluorescence and improve signal detection, all images were processed with rolling-ball background subtraction (radius = 50 pixels) prior to quantification. For each marker, individual fluorescence channels were separated and thresholded independently to determine the area of positive signal. Regions of interest (ROIs) were manually defined to correspond with the anatomical regions used for proteomic analysis, and measurements were collected within these defined ROIs. The positive area for each marker was quantified as the total thresholded area within the ROI and normalized to the ROI’s total tissue area (positive area/total area) to account for variability in tissue size across sections. Quantified data were analyzed using GraphPad Prism software. Outliers were identified using the ROUT method (Q=2%), and statistical significance between experimental groups was assessed using a Welch’s t-test; p<0.05 was considered statistically significant.

## Results

### Region-specific sex differences

Multiple distinct regions of human male (N = 7-12) and female (N = 5-8) lung were assessed using LCM-MS proteomic analysis to investigate sex differences across a spectrum of health and disease (Figure 1A). In control lungs, we assessed both airway and alveolar regions, while in IPF lung we investigated uninvolved distal airway, honeycombing airway, morphologically normal alveoli, aberrant basaloid epithelial cells, and fibroblastic foci (Figure 1B). PCA analysis of the resulting dataset demonstrated that proteome variation was driven more strongly by region than by sex, validating the utility of a tissue microenvironmental approach. Of all the regions, fibroblastic foci were the most divergent (Figure 1C). Two-way ANOVA corroborated these findings, with more differences observed due to region alone than to sex alone, though interestingly there were also significant interaction effects between the two variables, representing proteins changed by interdependent effects of sex on distinct regions (Figure 1D). Shared proteins across this analysis included factors involved in cilia function (RP2, DNALI1) and cytoskeletal changes during migration (SLK, DOCK9) (44–47). Several of these proteins are already known to be sex-linked (RP2, DNALI1, TSNAX), though their role in sex-specific lung biology has not been elucidated (44, 48, 49).

**Figure 1.**
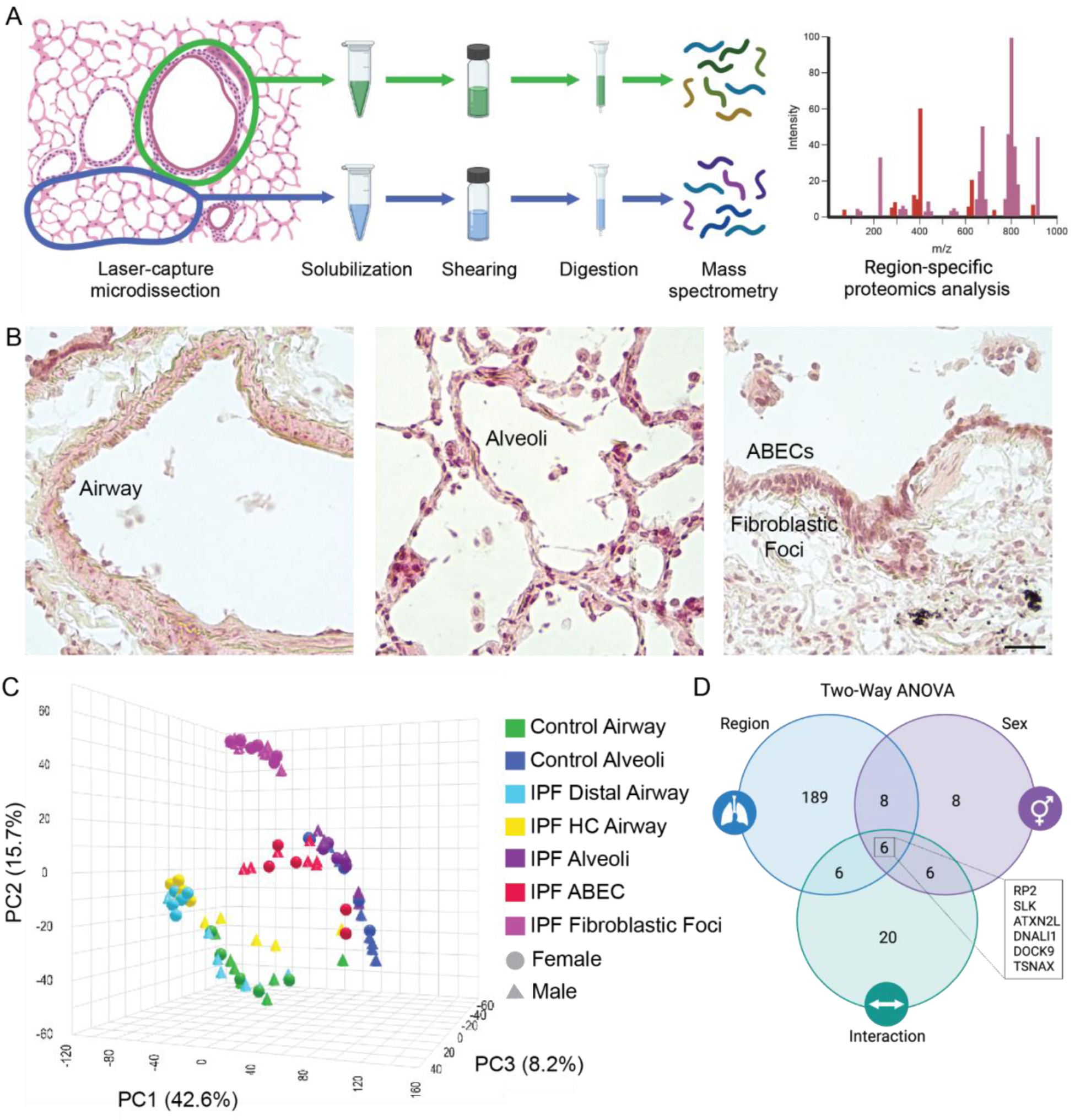
Region-specific sex differences in human lung. A) Experimental overview of region-specific proteomics analysis. B) Representative images showing airway, alveoli, and fibroblastic foci lined with ABECs. Scale = 50 µm C) PCA analysis of LCM-MS data annotated by region and sex reveals region-to-region proteomic separation. D) Two-way ANOVA analysis reveals differences due to both region and sex, as well as significant interaction between variables.

### Sex differences in control tissue

Regional sex differences in human lung were assessed by pathway enrichment analysis of differentially expressed proteins and PLS-DA VIP analysis to assess primary drivers of sex-biased differences (Figure S1, S2). Control alveoli show sex-biased expression of 321 proteins, including prominent male-biased expression of DLEC1 (Figure 2A). DLEC1 is a known tumor suppressor in lung and other cancers but also plays a role in spermatogenesis and cilia formation (50). The primary enriched pathway among these proteins is “Supramolecular Fiber Organization”, due to the presence of intermediate filament proteins (KRT6C and DES) and their interaction partners (DYNLRB1 and CNN3) (Figure 2B). Other enriched pathways include “Cytoskeleton in Muscle Cells” and “Vesicle-mediated Transport”, driven by differential expression of proteins like DYNLRB1 (Figure 2C) and SNAP29 (Figure 2D) which are involved in intracellular trafficking and part of downstream responses to classic inflammatory cytokines such as TNFα and TGFβ (51, 52). Of the proteins most involved in driving sex-biased differences in the proteome, as measured by VIP analysis, one notable protein is CNN3 (Figure 2E), which is overexpressed in both female alveoli and airway (FC=2.5) and associates with the wildtype MUC5B promoter (36). While all the CNN family members are involved in actin cytoskeleton dynamics, CNN1 has been implicated in pathological activation of both epithelial and mesenchymal cells in the context of IPF (53–55). In contrast, CNN3 has most commonly been studied in developmental contexts, but may also play a protective role in epithelial cells by negatively regulating fibrogenic responses (53, 56). Another protein of interest is STAT5A (Figure 2F), which is more frequently detected in male alveoli than in female. In addition to playing critical roles in inflammatory response and lung cancer progression, STAT5A has been implicated in driving sex differences in pulmonary hypertension and also associates with the most common gain-of-function MUC5B promoter variant driving IPF (36, 57, 58).

**Figure 2:**
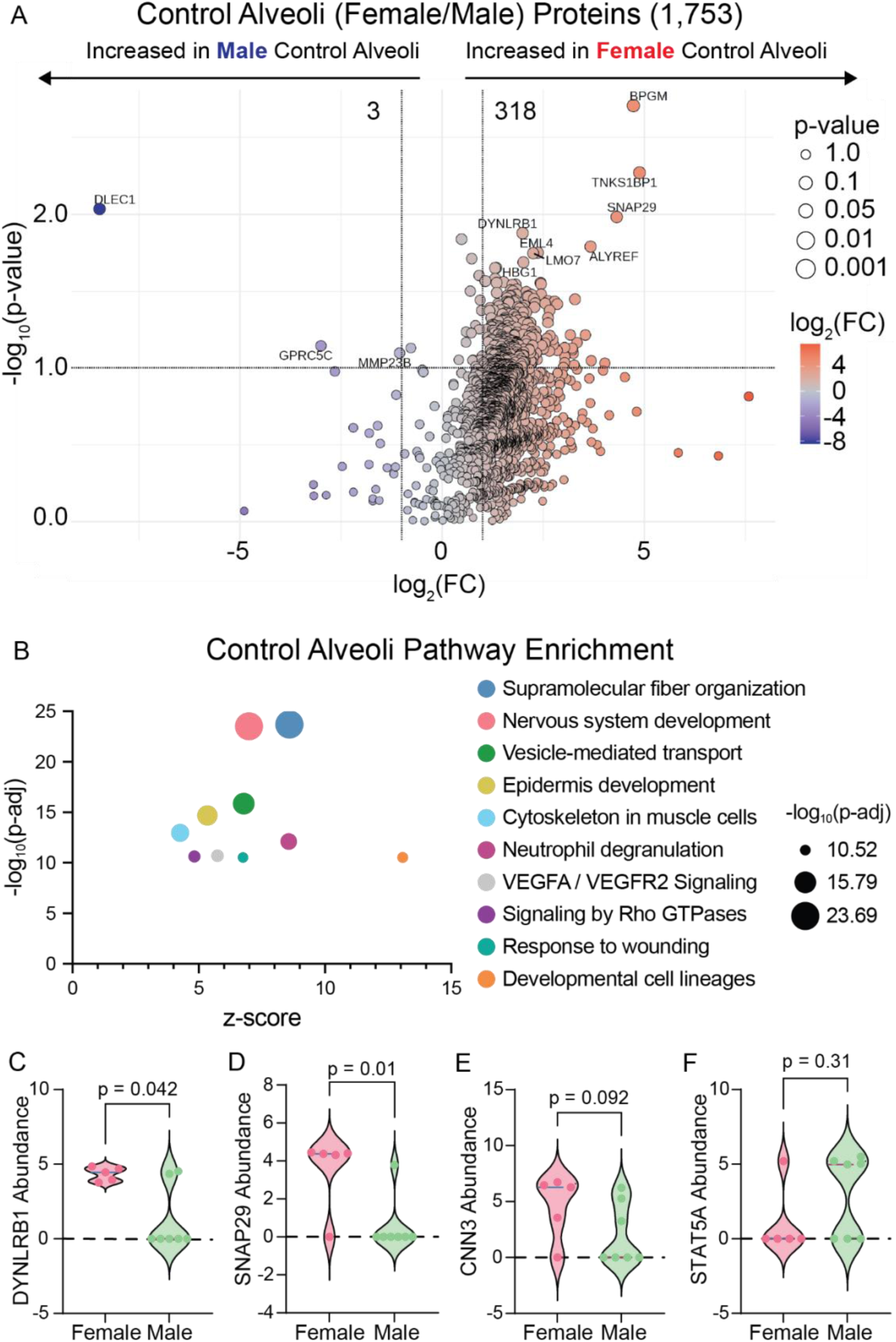
Sex differences in control alveoli. A) Volcano plot of differentially expressed proteins shows a trend towards protein overexpression in female alveoli. B) Pathway analysis shows prominent changes in pathways associated with cytoskeletal dynamics and transport. Protein abundance plots of C) DYNLRB1, D) SNAP29, E) CNN3, and F) STAT5A show expression changes in proteins driving overall sex differences.

Immunofluorescence staining confirmed the presence of both STAT5A and CNN3 in control alveoli, though staining quantification was not sensitive enough to detect differential expression by sex (Figure S3). Taken together, these results suggest baseline sex differences in the normal lung proteome might contribute to sex-biased epidemiology of lung diseases.

Control airway results showed 33 differentially expressed proteins, with more balance between female-skewed (14) and male-skewed (19) expression (Figure 3A). Among these proteins, notable hits include KLK5 (Figure 3B), an ECM protease involved in airway injury repair in the context of COPD, and SPINK5 (Figure 3C), a protease inhibitor that may modulate inflammatory responses in the context of asthma (59, 60). These proteolytic mediators drive enrichment of the primary differentially regulated pathways “Formation of Cornified Envelope” and the related “Differentiation of Keratinocytes” (Figure 3D). Another key protein driving sex differences is LRP1, which is overexpressed in male airway (FC=2.65) (Figure 3E). LRP1 is involved in the formation of endocytic vesicles, and thus recycling of extracellular proteases, but is also implicated in regulating fibroblast activation and ECM synthesis (61–63). Relatedly, female airways show higher expression of the critical ECM molecules COL1A1 (FC=1.08) and ELN (FC=1.90), the latter of which was also detectably differentially expressed by staining (Figure 3F). In line with these observed ECM changes, studies have suggested that there may be sex differences in overall lung compliance and elasticity, though fully understanding these effects is complicated by variables including height and age (64–66). Still, such underlying differences between the normal lung proteome of male and female patients could be a basis for both differences in lung mechanics and sex-biased epidemiology of various lung diseases.

**Figure 3:**
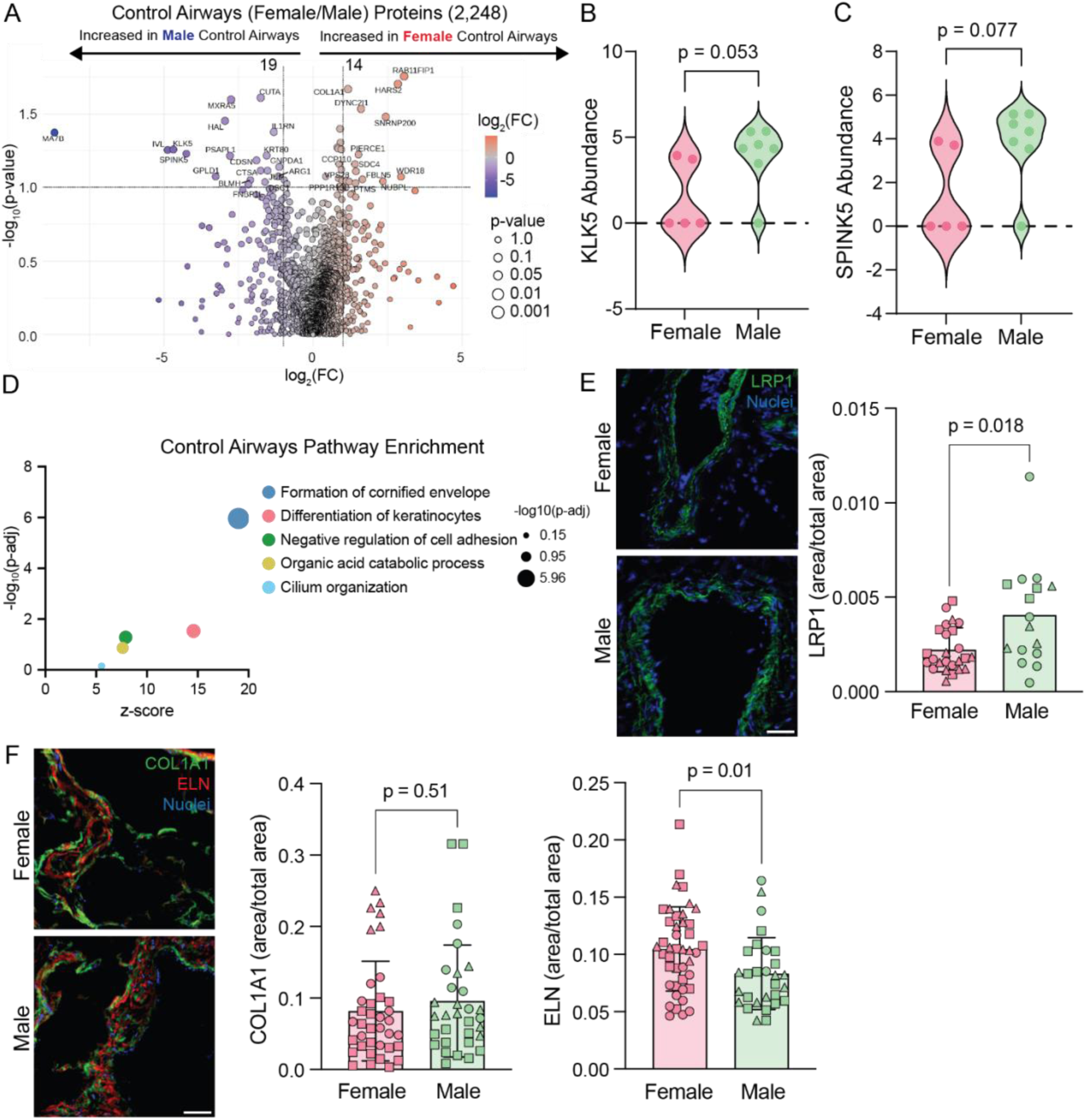
Sex differences in control airway. A) Volcano plot of differentially expressed proteins. Protein abundance plots of B) KLK5 and C) SPINK5 show male-biased expression of proteolytic mediators. D) Pathway analysis includes pathways with strong protease presence. E) Immunostaining analysis confirms LRP1 overexpression in male airways. F) Staining for COL1A1 and ELN shows increased ELN in female airways. Scale bars = 50 µm.

### Sex differences in IPF airways

Sex differences become much more pronounced in IPF lung, particularly in airways (Figure S4). Distal airways show differential expression of 800 proteins (Figure 4A), while honeycombing airways have 1,679 differentially expressed factors (Figure 4B), with both regions showing a skew towards protein overexpression in female lung. Distal airways show sex differences in pathways related to intracellular trafficking and cytoskeletal dynamics (Figure 4C), including “Neutrophil Degranulation” and “Intracellular Protein Transport,” which are also present in honeycombing airways, along with additional pathways related to cellular metabolism (Figure 4D). In both types of airway regions, sex-biased expression of ECM molecules is even more exaggerated than in control airway, with prominent female-biased overexpression of COL1A1 and ELN detectable by staining (Figure 4E). COL1A1 is a classic marker of IPF, largely due to increased expression in progressively expanding fibroblastic foci, but it may also contribute to disease progression from the airway (67, 68). In contrast, ELN is known to increase in the airway of fibrotic lung, and its role in fibrotic progression is more likely reliant on architecture and crosslinking rather than simple quantity (69–71). In line with these other ECM molecules, a driver of airway proteomic differences is COL6A6, which is overexpressed in a subset of female airways as demonstrated by staining (Figure 4F). Collagen VI is a critical structural component of the basement membrane that helps regulate lung elasticity and may drive progression of pulmonary fibrosis (72, 73). In contrast to the female-biased expression of these structural ECM components, male airways show increased expression of the PRSS family of serine proteases (Figure 4A-B), suggesting enhanced ECM degradation, all of which suggests an enhanced airway-centered response to IPF in female patients.

**Figure 4:**
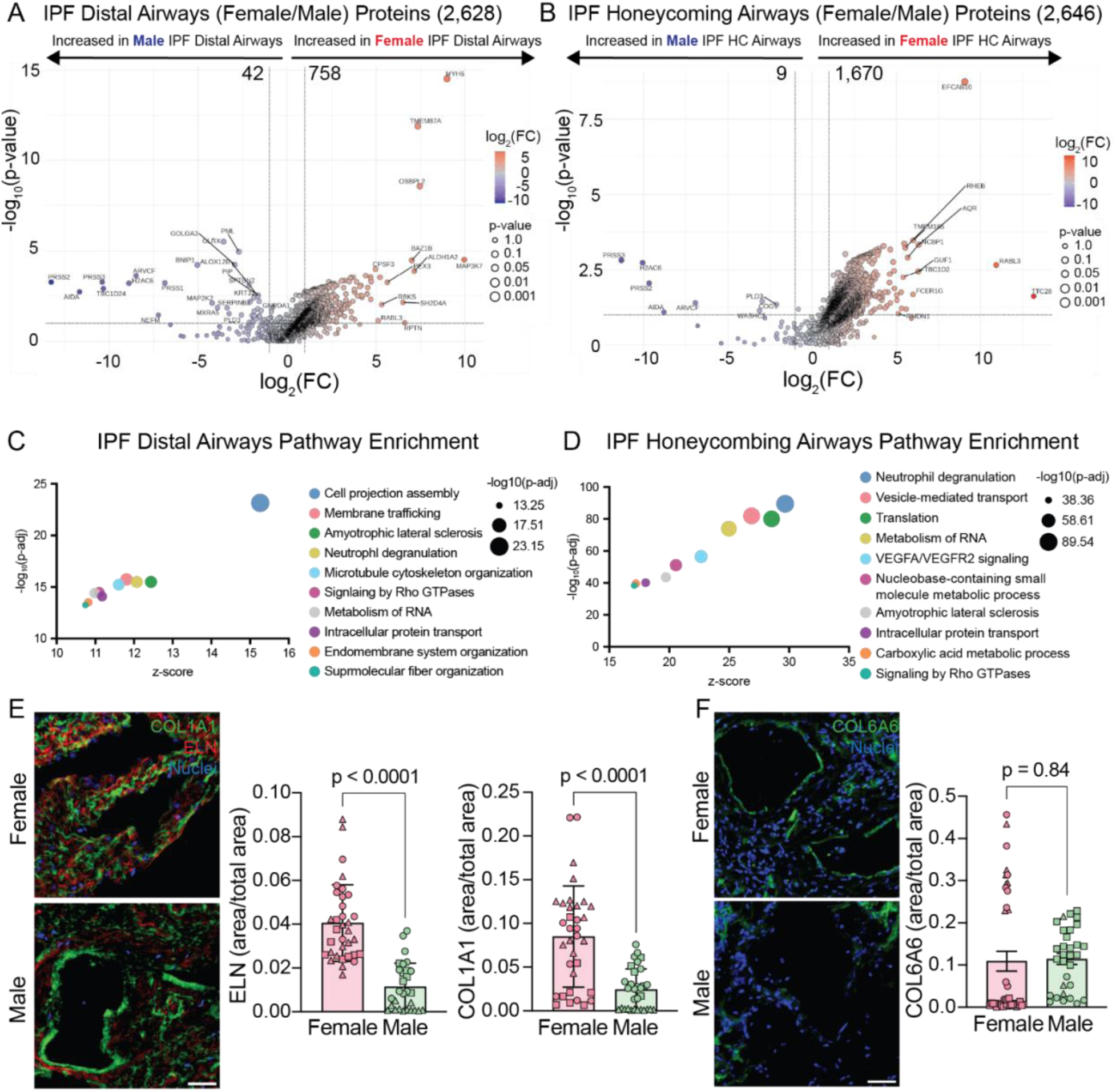
Sex differences in IPF airway. Volcano plots of differentially expressed proteins in A) distal and B) honeycombing airways in IPF lung reveal pronounced sex differences. Pathway analysis in C) distal and D) honeycombing airways show changes in cytoskeletal dynamics and cellular metabolism. E) Immunostaining for COL1A1 and ELN shows increased expression in female airways. F) Immunostaining for COL6A6 shows diversity of expression particularly in female airways. Scale bars = 50 µm.

### Sex differences in IPF distal lung

Morphologically normal alveoli from IPF patients show 82 differentially expressed proteins between sexes (Figure 5A), with the top represented pathways being “Neutrophil Degranulation” and “Cornified Envelope Formation” (Figure 5B). As with results for control alveoli, these pathways were enriched largely due to the male-biased expression of protein trafficking factors including VAMP8, KRT12, and related proteins. Other differentially expressed pathways included “Peptide Crosslinking” and “Degradation of the Extracellular Matrix,” which could relate to changes in tissue mechanical properties underlying the spread of disease.

**Figure 5:**
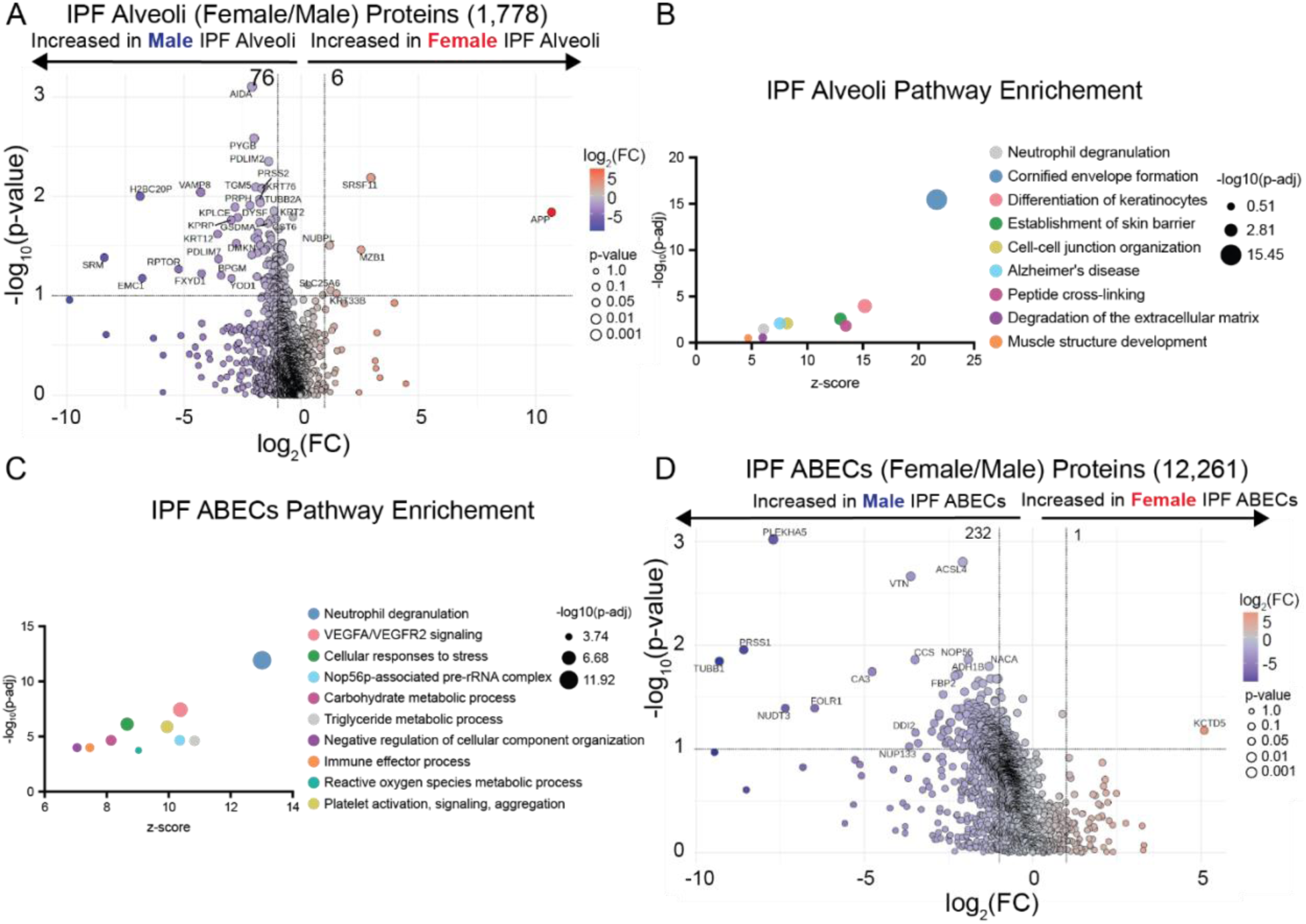
Sex differences in IPF epithelial regions. A) Volcano plot of differentially expressed proteins in morphologically normal alveoli from IPF patients shows protein expression skewed towards males. B) Pathway enrichment analysis shows changes in pathways involving proteolysis and protein crosslinking in IPF alveoli. C) Pathway enrichment analysis shows changes in pathways related to cellular stress and metabolism in ABECs. D) Volcano plot of differentially expressed proteins in ABECs shows protein expression skewed towards males.

In diseased regions, aberrant basaloid epithelial cells (ABECs) showed differences in pathways related to cellular stress and metabolism (Figure 5C), with male-biased expression of 232 proteins including VTN (Figure 5D, S5), an ECM component implicated in cellular differentiation and control of apoptosis during fibrotic development (74). One strong driver of sex differences was AHNAK2, which trends towards being higher in male ABECs (FC 2.24). AHNAK2 was also detectable by staining, with a slight trend towards overexpression in male ABECs (Figure S5).

AHNAK2 has been studied in the context of lung cancer, where it promotes epithelial-to-mesenchymal (EMT) transition of lung adenocarcinoma cells, but it is also upregulated in IPF, where it appears to play a similar role in TGFβ-induced EMT (75–77). A heightened EMT phenotype in male ABECs is consistent with increased cellular metabolism and stress markers and could play a role in worsening the progression of underlying fibroblastic foci. Taken together, these data suggest greater levels of distal epithelial cell dysfunction in male IPF.

Meanwhile, fibroblastic foci themselves show a strong skew towards elevated protein expression in females with 107 proteins overexpressed in females (Figure 6A). Many of the top differentially regulated pathways were involved in cellular interaction with the extracellular environment, including multiple pathways related to cytoskeletal dynamics, cellular adhesion, and ECM organization (Figure 6B). Strong drivers of sex differences included minor collagens like COL11A1 (Figure 6C) and COL8A2 (Figure 6D), as well as basement membrane components like LAMB3, which was detectably higher in female fibroblastic foci by staining (Figure 6E). LAMB3 is upregulated in models of injury-induced pulmonary fibrosis but may actually play a protective role in facilitating healthy re-epithelialization rather than driving epithelial cell injury phenotypes (78–80). Overall, these data potentially suggest a greater ECM diversity in female foci, including factors that might mediate fibrotic response, a phenomenon which calls for more detailed study.

**Figure 6:**
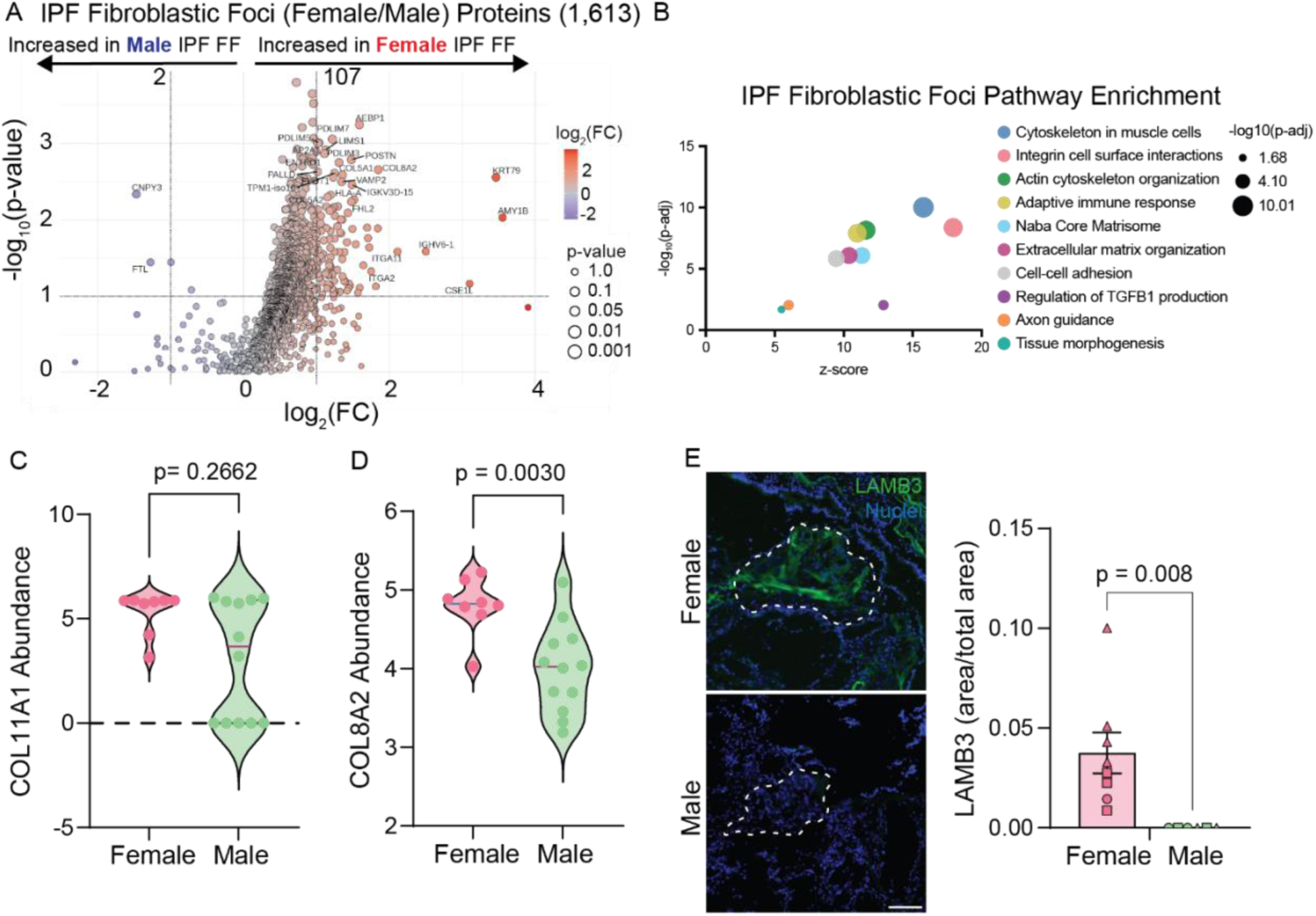
Sex differences in fibroblastic foci. A) Volcano plot of proteins from fibroblastic foci shows skew towards protein overexpression in female samples. B) Pathway analysis shows changes in cytoskeleton, cellular adhesion, and ECM organization pathways. Protein abundance of C) COL11A1 and D) COL8A2 show consistently high expression in female samples. E) LAMB3 immunostaining shows overexpression in female foci. Scale = 100 µm.

### Conclusions

Proteomic studies have revealed a valuable snapshot of diseased lung, and uncovered biomarkers of IPF that are upregulated during the progression of disease (81), but further studies are required to elucidate whether upregulated factors are playing contributing roles to disease, or protective roles against progression (82–84). In this study, we demonstrated that human lungs show baseline sex-differences in protein expression, with more pronounced differences in airway but also sex-bias in the alveolar proteome. Across all regions in both control and IPF lung, one common sex-biased pathway was “Neutrophil Degranulation,” a cellular process which involves formation of vesicles (granules) to be trafficked along the cytoskeleton and fused with the plasma membrane for exocytosis (85). Extracellular vesicles accumulate during IPF and mediate cell–cell crosstalk, suggesting that lung-wide sex differences in this pathway could profoundly influence disease progression (86, 87). IPF lungs tended to show sex-biased enrichment of metabolic pathways across all regions, while distal lung regions specifically showed enrichment of pathways related to cell adhesion and ECM organization, critical contributors to disease progression.

Overall, sex-specific protein expression may begin to explain why similar risk factors (e.g. smoking) lead to different disease pathology in different patients (COPD in women, IPF in men). Sex-differences in diseased lungs are more pronounced, with female patients showing pronounced ECM expression in airways and ECM diversity in fibroblastic foci, while male patients exhibit increased markers of epithelial cell dysfunction. Reduced survival of men with IPF may be due to the engagement of such sex-specific profibrotic pathways. The sex differences observed here also have potential implications for treatment response, due to sex-biased regulation of several critical drug-target pathways. VEGF signaling, a target of the classical antifibrotic drugs nintedanib and pirfenidone, showed sex-biased regulation in control alveoli, honeycombing airway, and ABECs (88, 89). Similarly, Rho-GTPase activity, a pathway linked to the mechanism of action of the recently-approved therapeutic nerandomilast, was differentially regulated in control alveoli, IPF distal airway, and honeycombing airway (90). In fibrotic foci, sex-biased regulation of TGFβ production has implications for response to pirfenidone treatment (91). Taken together, these results highlight that considering sex as a biological factor in both models of chronic lung disease and in clinical settings is essential.

## Acknowledgements

Human lung samples were provided by the Human Tissue and Cell Core at the University of Colorado Anschutz, supported by NIH-NHLBI grant P01HL162607. Mass Spectrometry was performed by the Mass Spectrometry Proteomics Shared Resource Facility [RRID SCR_021988] at the University of Colorado Anschutz.

This work was supported by funding from the National Heart, Lung, and Blood Institute of the National Institutes of Health (NIH) under awards R01 HL153096 (CMM, RB, HN, MCM), P01-HL162607 (JAH, DAS), and R01-HL097163, R01-HL148437, R01-HL158668, P01-HL092870, UH3-HL123442 and UG3-HL151865 (DAS). Additional funding came from the NIH/NCATS Colorado CTSA Grant Number T32 TR004367, UM1 TR004399 (HN) and the VA Merit Review (IO1BX005295, DAS).

## Data Availability

Proteomic data are available under the identifier PXD058626 at the ProteomeXchange Consortium (https://proteomecentral.proteomexchange.org/).

Additional data required to reproduce this work are available at doi: 10.17632/jtf74kncvv.1.

