## Supplemental Material for "Proteomic analysis reveals regional sex differences in healthy and fibrotic human lung"

### Supplementary Material

Table S1: Subject Demographics

| Disease state | Sex | Age | MUC5B genotype |
|---|---|---|---|
| Control | Female | 49 | GT |
| Control | Female | 61 | GG |
| Control | Female | 62 | GG |
| Control | Female | 67 | GT |
| Control | Female | 79 | GG |
| Control | Male | 49 | GT |
| Control | Male | 57 | GG |
| Control | Male | 63 | GT |
| Control | Male | 66 | GT |
| Control | Male | 69 | GG |
| Control | Male | 71 | GG |
| Control | Male | 74 | GT |
| IPF | Female | 59 | GG |
| IPF | Female | 68 | GG |
| IPF | Female | 68 | GT |
| IPF | Female | 69 | GG |
| IPF | Female | 69 | GT |
| IPF | Female | 70 | GT |
| IPF | Female | 71 | GT |
| IPF | Female | 76 | GT |
| IPF | Male | 47 | GT |
| IPF | Male | 52 | GG |
| IPF | Male | 65 | GG |
| IPF | Male | 65 | GT |
| IPF | Male | 66 | GG |
| IPF | Male | 66 | GT |
| IPF | Male | 67 | GT |
| IPF | Male | 67 | TT |
| IPF | Male | 68 | GG |
| IPF | Male | 68 | GT |
| IPF | Male | 68 | GT |
| IPF | Male | 73 | GT |

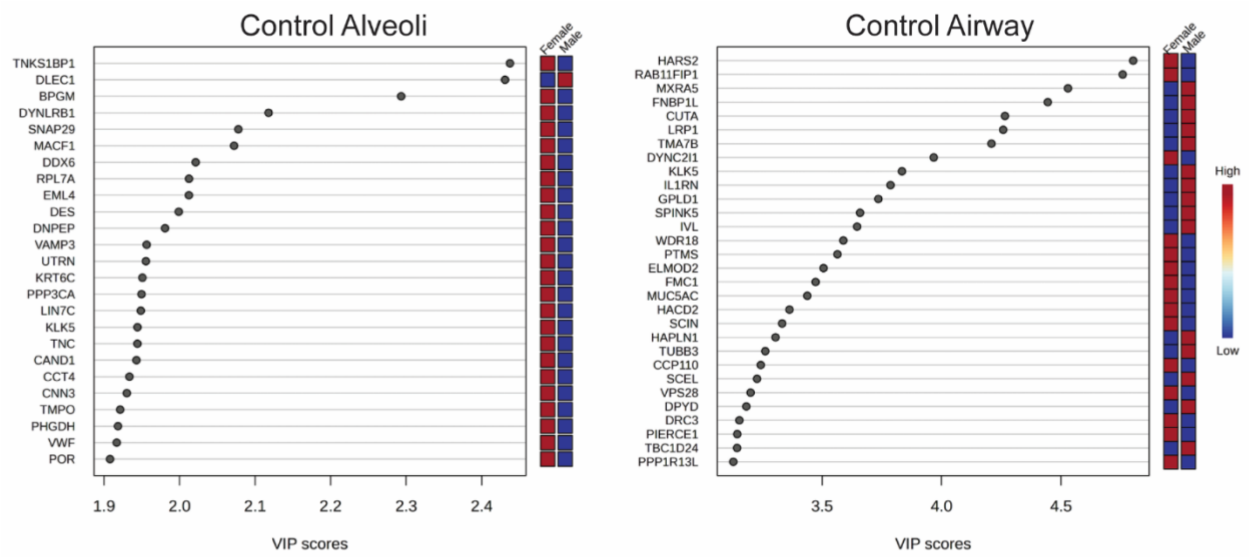


Figure S1: Variable importance in projection plots show proteins driving sex differences in control lung.

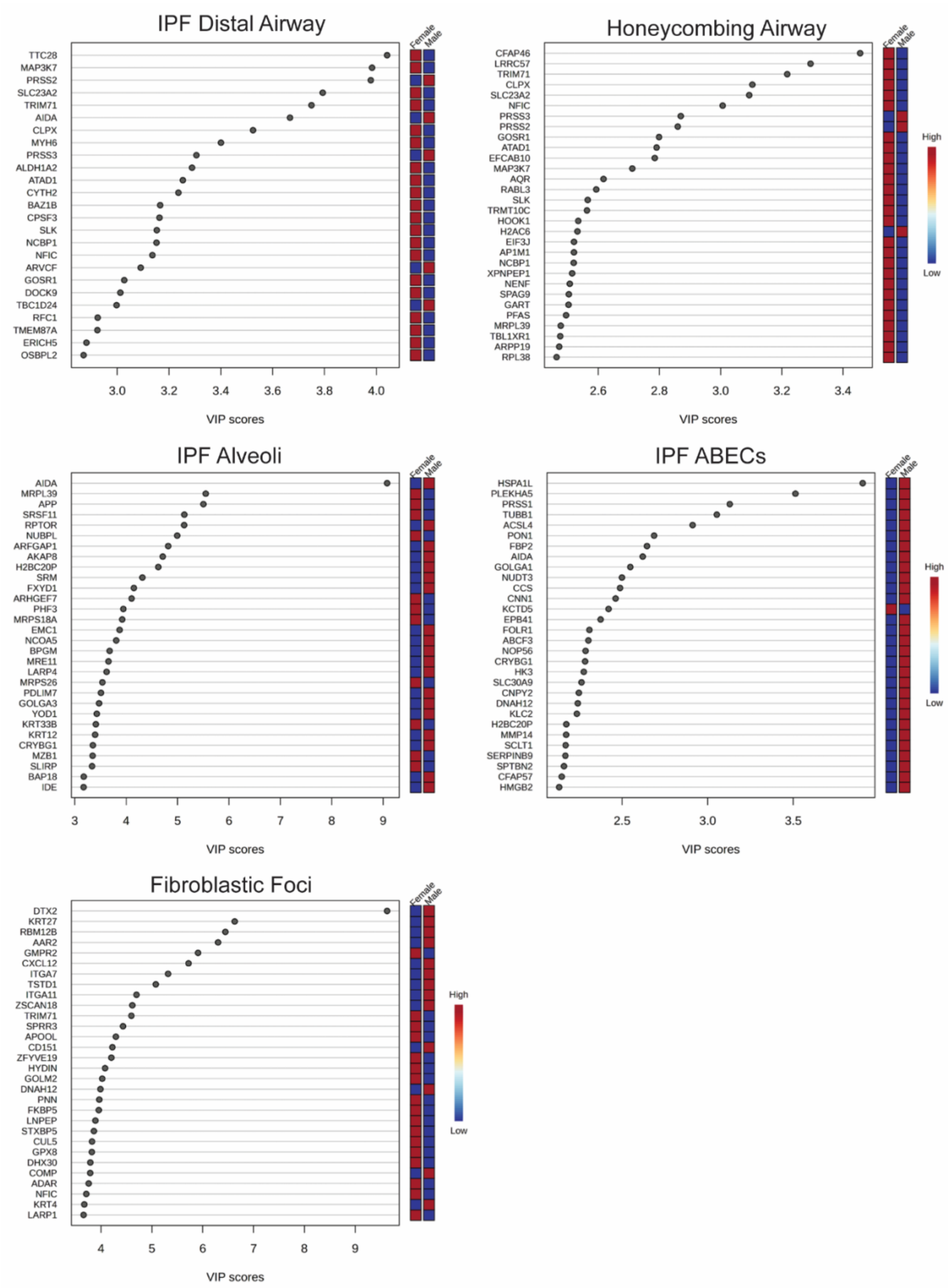


Figure S2: Variable importance in projection plots show proteins driving sex differences in IPF lung.

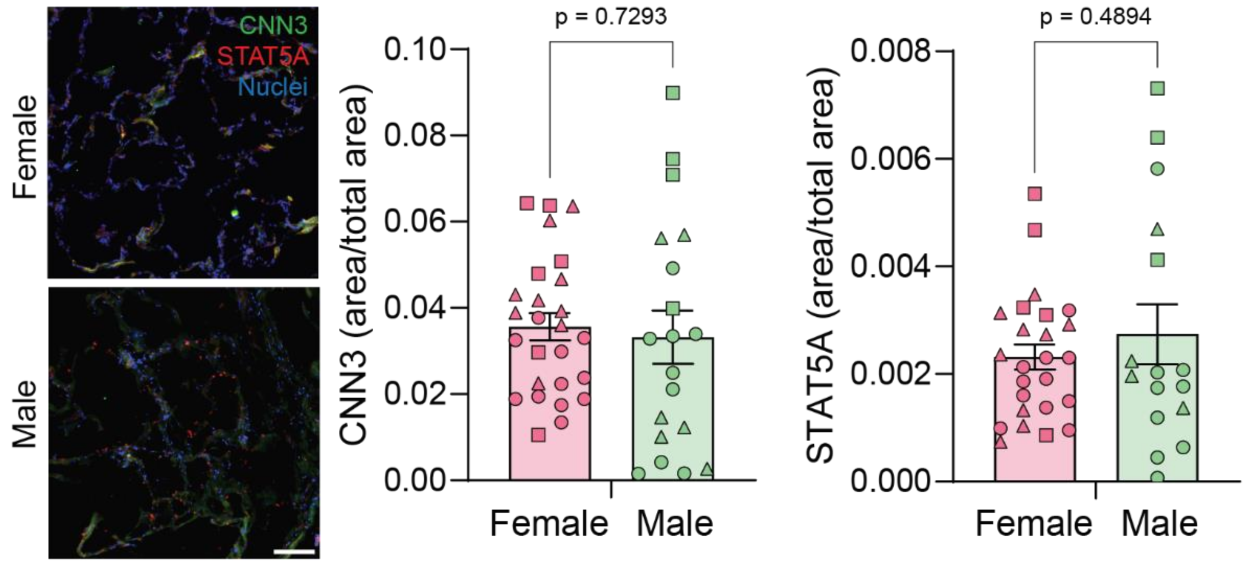


Figure S3: Immunostaining shows CNN3 and STAT5A both detectable in human control alveoli. Scale = 100 μm

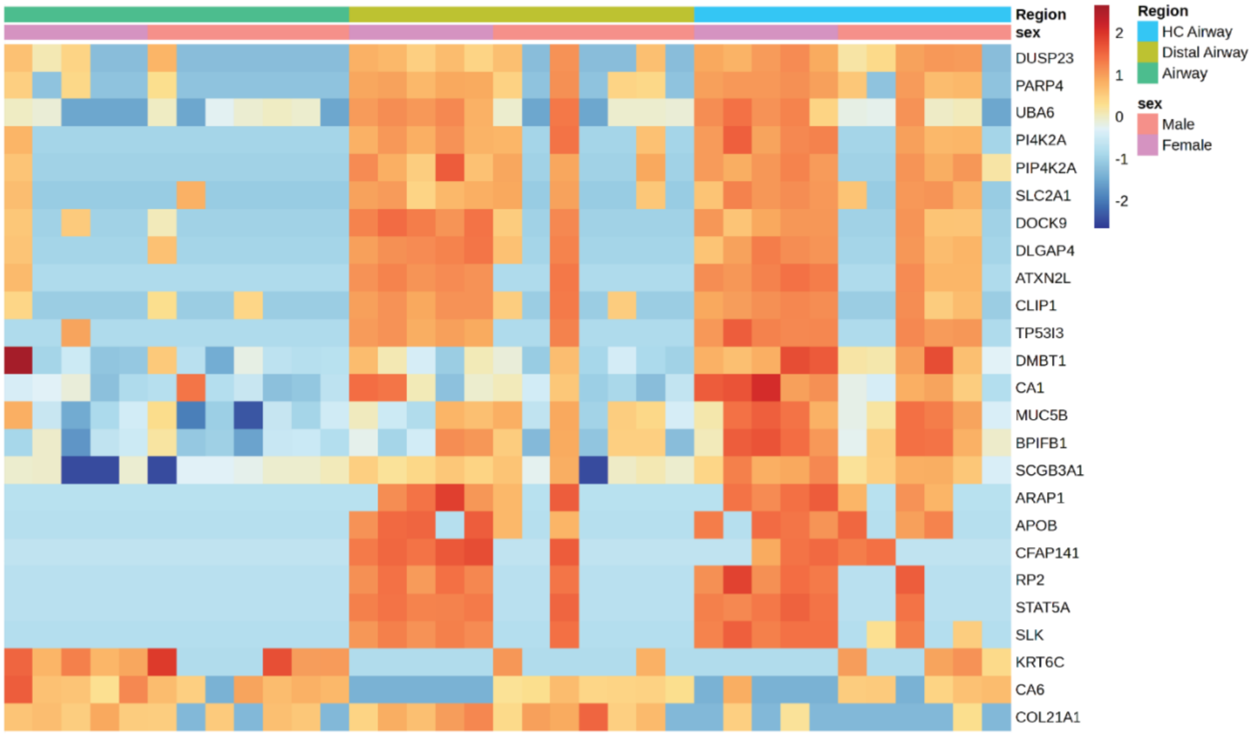


Figure S4: Heatmap of sex-biased proteins in control airway, IPF distal airway, and honeycombing airway shows that disease exacerbates sex differences.

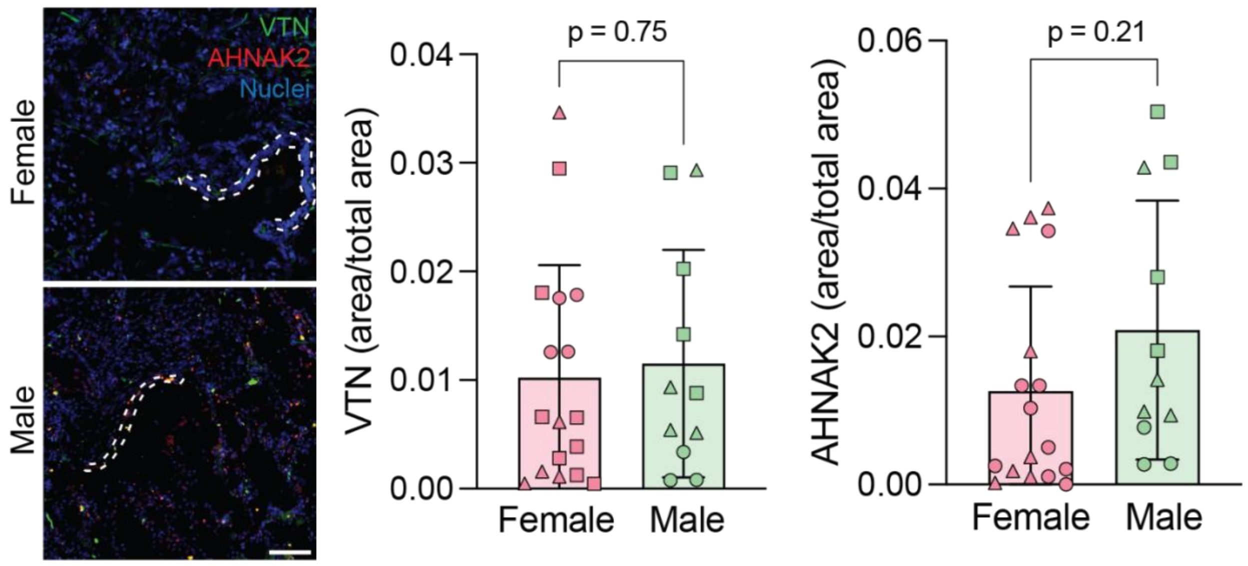


Figure S5: Immunostaining shows VTN and AHNAK2 both detectable in IPF ABECs. Scale = 100 µm
